# Ontogenetic expansion and regionalization of the triatomine compound eye supports flight-related vision

**DOI:** 10.64898/2026.04.12.717996

**Authors:** Tomás Manuel Chialina, Hernán Gustavo Gentilli, Sebastián A. Minoli, Martín Berón de Astrada

## Abstract

Triatomines are the vectors of Chagas disease, one of the main endemic diseases from South to North America, now expanding to other continents. These hemimetabolous insects have been considered poorly visual animals. However, recent findings challenge this idea.

Here, we used *Rhodnius prolixus* as a model species to comprehensively characterize triatomine compound eyes. We found that in the adult stage, eye size significantly exceeds the dimensions predicted by the nymphal eye growth rate. Moreover, while the compound eye grows symmetrically in its dorsal and ventral directions throughout the nymphal instars, in the adult, the eye undergoes greater ventral growth, resulting in a dorsoventrally asymmetrical eye.

By studying a bright pseudopupil induced by fluorescence in natural mutant animals, we observed no major differences in sampling resolution between the last nymphal instar and the adult stage. However, the adult eye possesses significantly larger ommatidia, particularly in its ventral region, shifting the area of highest sensitivity from the equatorial region in the nymphal instars to the ventral region in the adult.

A similar eye growth pattern was observed in *Triatoma infestans* and *Panstrongylus megistus*. The analysis of photographic records from 39 species across 10 genera indicates that an asymmetrical eye is the predominant eye pattern in adult triatomines. Notable exceptions in wingless adults of *Mepraia spinolai*, reveal a tight association between possessing a large asymmetrical eye and the presence of wings. This suggests that vision might support triatomine dispersal flights among other visual behaviors.

**Significance Statement:** Kissing bugs are hematophagous insects known for being the vectors of Chagas disease, one of the main endemic diseases in the Americas. Vision was not considered a relevant sensory system in these insects. Here, we show that their eyes increase in size beyond expected by ontogeny and become asymmetrical when transitioning from the last nymphal instar to the adult stage. The eyes undergo a ventral expansion that shifts the region of greatest light sensitivity from the equatorial zone in nymphs to the ventral region in adults. We found this asymmetrical eye only in winged kissing bugs, suggesting that vision supports flight. This is relevant in ecological and epidemiological terms since kissing bugs disperse by flight for habitat colonization and host-seeking.

## Introduction

Triatomines (Heteroptera: Reduviidae: Triatominae), commonly known as kissing bugs, are a particular subfamily of reduviid bugs whose principal diagnostic character is their obligatory hematophagy (1). Kissing bugs feed on vertebrate blood throughout their life cycle, a condition likely derived from the primitive predatory habits of the Reduviidae family (1, 2). Triatomines are potential vectors of the parasite *Trypanosoma cruzi*, which they can transmit to their mammalian hosts, causing Chagas disease, a condition that affects thousands of people each year in the Americas (3). This is one of the main reasons that have motivated extensive studies of the ecology, physiology, and behavior of kissing bugs (e.g. (4–6)).

Over 150 species of triatomines, grouped into 17 extant genera (7), inhabit diverse wild and domestic microhabitats. These habitats range from vertebrate shelters (subterranean or arboreal), bird nests, and human dwellings to cattle corrals and chicken coops. They also include spaces with no direct relationship to the hosts, such as rock outcrops, trees, palm crowns, various plants, and epiphytes (8, 9). Kissing bugs may live and feed inside the nests or shelters of their vertebrate hosts (sit-and-wait strategy, nest specialists), or they may actively search for and opportunistically feed on diverse hosts that temporarily share their habitat (stalker strategy, opportunistic host generalists) (10). Unlike nymphs, winged adults can disperse by flight, driven by hunger, high population densities or other stimuli (11, 12). In these situations, they can actively search for hosts, regardless of whether they previously lived as “sit-and-wait” or “stalkers” foragers.

Probably due to their crepuscular and nocturnal habits (13, 14), triatomines have not been considered highly visual insects. Studies on their visually-guided behaviors are rather scant compared to studies on other sensory modalities (6). It was observed that several species avoid illuminated spaces guided by a marked negative phototaxis (15–17). However, some triatomine species also show positive phototaxis to dim punctual light sources, as evidenced in both laboratory experiments and natural environments (18–22). In addition, reports of arrivals of flying triatomines to house lights are frequent in endemic areas (23, 24). While these phototactic behaviors are rather simple in their demands of visual system capabilities, visual behaviors that involve motion or object detection require more complex visual processing (e.g. (25)). With this respect, Lazzari and Varjú (26) found that *T. infestans* can fixate moving targets in the posterior-lateral region of their visual field. The authors suggested that this capability would allow the animals to visually track an eventual predator while still being able to escape in a direction away from it. More recently, Chialina et al. (27) found that *R. prolixus* consistently responds to moving visual danger stimuli by either freezing or directionally escaping. Moreover, they observed that *R. prolixus* can rapidly switch between these behaviors based on ongoing stimulus information in an adaptive manner, similar to what has been observed in highly visual arthropods (e.g., locusts (28), crabs (29), and flies (30)).

The visual system supporting spatial vision in reduviids, and particularly in triatomines, has been scarcely characterized (but see (22, 31–34)). Triatomines count with a pair of rather globose compound eyes laterally protruding from their head capsule (35). These are apposition compound eyes in which the ommatidia possess open rhabdoms. The rhabdom is composed of a ring of six rhabdomeres (R1-6) surrounding a central pair of rhabdomeres (R7-8) (33, 36). The only two species of this group in which quantitative analyses of the eyes have been carried out, *Triatoma infestans* and *Rhodnius prolixus*, possess between 30 and 400 ommatidia depending on the developmental stage (37). These eyes exhibit features of eyes capable of adapting to strong variations in ambient light intensity. For instance, it has been observed that screening pigments inside retinular cells and primary pigment cells form a dynamic ‘pupil’ in the ommatidia of *R. prolixus* and *T. infestans*, whose diameter changes with light intensity or both light intensity and time of the day (31, 33, 38).

Although the neural architecture of the ganglia dedicated to processing visual information from triatomine compound eyes remains virtually unknown, Insausti (32) observed that the optic lobes of *T. infestans* are voluminous, and make up the majority of the protocerebrum, the largest region of the brain in these animals. Additionally, a recent comparative review by Hearn and Wolff (39) underscored the notable development of the optic lobes in the brain of kissing bugs with respect to other hematopahagous arthropods.

The epidemiological importance of triatomines, along with recent findings on neural investment in vision and the complex visual behaviors they exhibit, motivated us to comprehensively characterize the morphology of their compound eyes. Using *R. prolixus* as a model species, we conducted a detailed study of the eye across its different developmental stages. As hemimetabolous insects, the development of kissing bugs consists of five sequential nymphal instars followed by an adult stage. Although the anatomical changes are quite gradual and progressive across stages, adults distinctively develop reproductive maturity, two pairs of wings, two pairs of exocrine thoracic glands and one pair of ocelli (40, 41). In the current study, we found that throughout the nymphal instars, the compound eye grows symmetrically in both its dorsal and ventral directions. However, in the adult, the eye undergoes greater ventral expansion, resulting in an eye that is no longer dorsoventrally symmetrical. Furthermore, in the adult stage, eye size significantly exceeds the dimensions predicted by the nymphal eye growth rate. By studying a bright pseudopupil induced by fluorescence in natural mutant animals, we observed no evident differences in the visual fields of the eyes between the last nymphal instar and the adult stage. Similarly, sampling resolution presented no major differences along the studied transects of the eye. However, the adult eye exhibits larger ommatidia, particularly in its ventral region, shifting the area of highest sensitivity of the eye from the equatorial region in the nymphal instars to the ventral region in the adult. When we analyzed other triatomine species, such as *T. infestans* and *Panstrongylus megistus*, we observed a similar pattern of eye growth between the last nymphal instar and the adult. Furthermore, when we analyzed the available photographic records of the eyes of several other triatomine species - 39 species from 10 genera - we observed that in almost all cases, the adult eye extends more ventrally than dorsally, exhibiting the largest ommatidia in the ventral region. One important exception was found in wingless adults of *Mepraia spinolai*. In this particular case, we found that the compound eye remains relatively small and dorsoventrally symmetrical, contrasting with the large and dorsoventrally asymmetrical eye of the winged morph of the same species. This indicates a tight association between possessing a large asymmetrical eye and the presence of wings. This association strongly suggests that vision might support triatomine dispersal flights, along with other behaviors we discuss here.

## Results

### 3.1 Eye growth morphometrics

In many different species, vision serves as a key sensory input for guiding locomotor behaviors such as walking or flying. Like most triatomines, *R. prolixus* is capable of walking throughout all its stages, whereas flight is restricted to the adult stage (Fig. 1 A-C). Their compound eyes are consistently positioned laterally, presenting an approximately hemispherical shape protruding from the head. As in various triatomines, the head of *R. prolixus* is elongated along the anteroposterior axis. Consequently, the anterior and posterior regions of the head constitute a steric impediment to vision. The lateral protrusion of the compound eyes, therefore, facilitates an expanded visual field along this axis, particularly towards the anterior region (Fig. 1 D-F).

**Figure 1.**
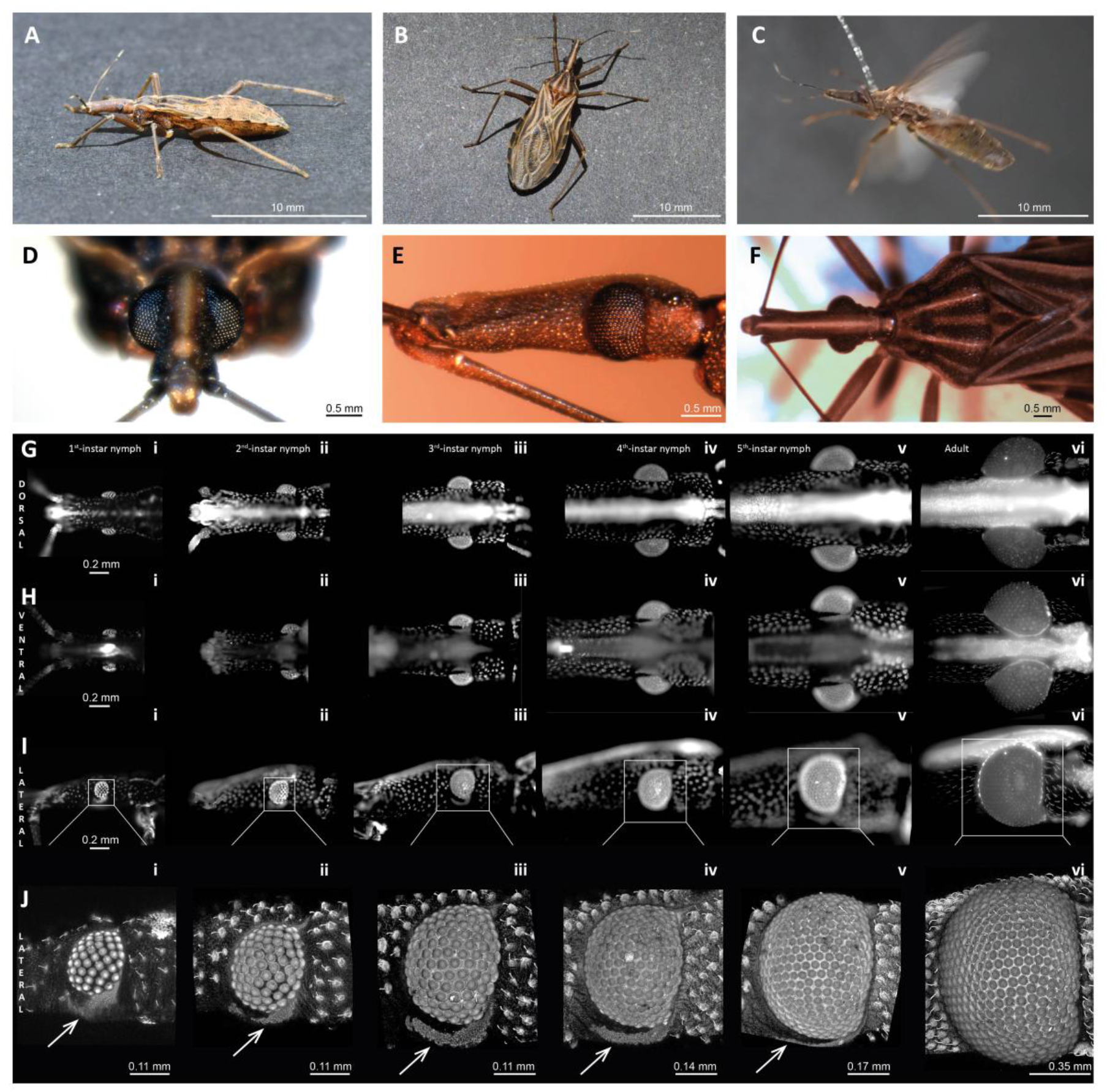
General morphology and post-embryonic growth of the compound eye in *R. prolixus*. **A)** Fifth-instar nymph in walking posture. **B-C)** Images illustrating the locomotion modes of an adult: walking and flight, respectively. **D)** Frontal, **E)** lateral and **F)** dorsal views of the head of an adult. **G)** Dorsal, **H)** ventral and **I)** lateral views of the heads across all developmental stages under an epifluorescence microscope, respectively. **J)** Confocal microscopy images of the compound eyes in lateral view. Each image consists of a Z-projection of the compound eye for all stages. For *G-J, i-v* represent the nymphal instars (I to V), and *vi* represents the adult stage. Arrows indicate a bright curved band ventral to the compound eye, present only in nymphal instars. Scale bars in *G-I* apply to all images (*i-vi*) within each respective row.

Figure 1 G-I displays the dorsal, ventral, and lateral views of *R. prolixus* head across its different instars. A cursory inspection of the images reveals that the eyes grow with head size, always with the characteristic lateral protrusion. However, there is a considerable shift in eye size in the adult stage, where the eye grows considerably more relative to head size compared to the nymphal instars (Fig. 1 G_vi_-I_vi_; Fig. 2). The size increment is more pronounced along the dorsoventral extent of the eye, significantly reducing the interocular distance in the dorsal and ventral regions of the head (Fig. 1 G_vi_-H_vi_), though the change is more dramatic in the latter (Fig. 1 H_vi_). Notably, a distinct bright mark shaped like a curved band is consistently observed on the dark cuticle ventral to the compound eye across all nymphal instars, occupying a constant position relative to the head (arrows in Fig. 1J_i-v_). This curved band is absent in the adult stage as the corresponding cuticular area is occupied by the eye (Fig. 1J_vi_).

**Figure 2.**
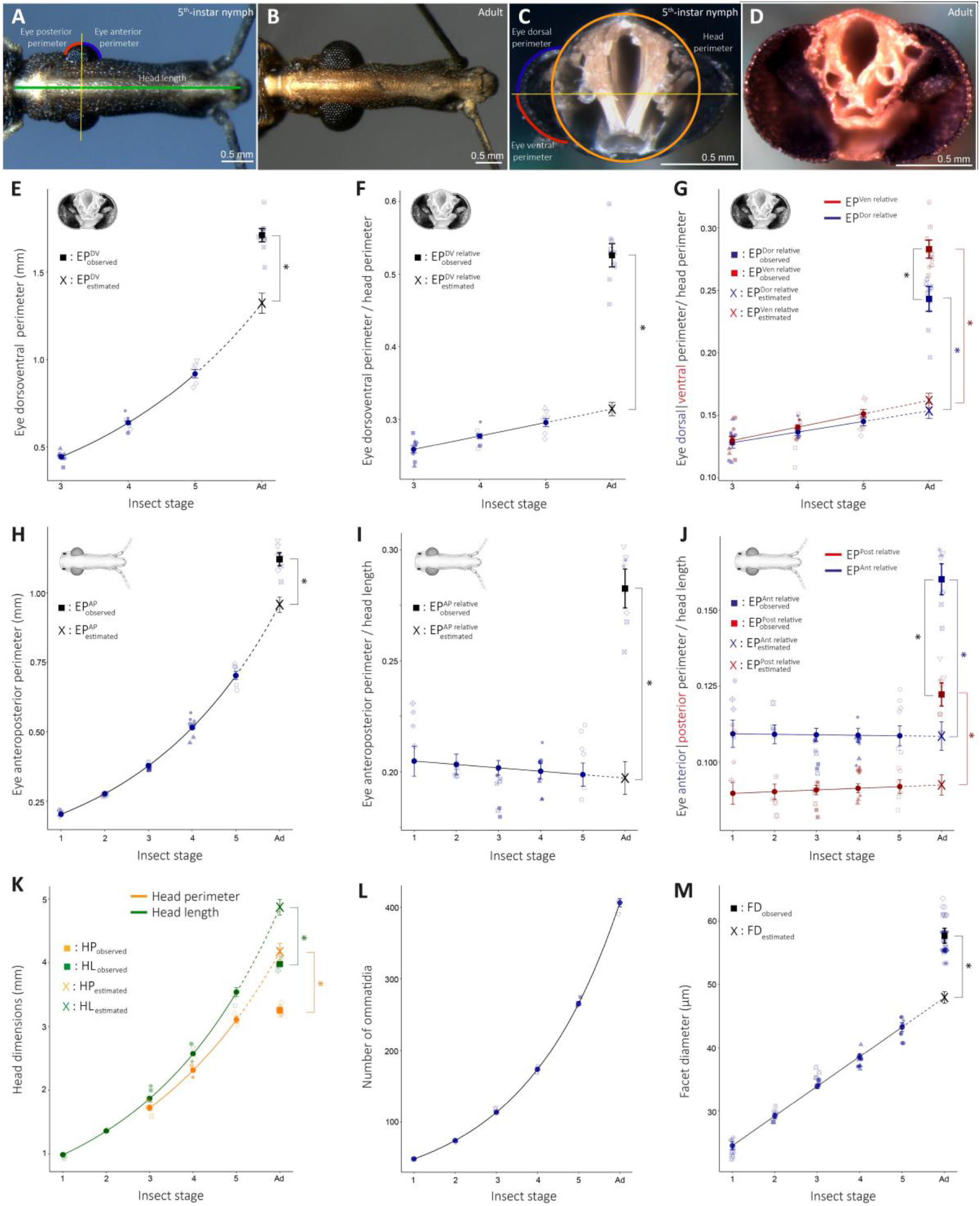
Quantification of eye growth of *R. prolixus*. **A-B)** Dorsal views of the heads of a fifth-instar nymph and an adult, respectively. **C-D)** Transversal sections of the heads of a fifth-instar nymph and an adult, respectively. *A* and *C* also illustrate the measurements performed in each view. **E-F)** Post-embryonic growth of the eye dorsoventral perimeter, expressed both in absolute terms (*EP*^*DV*^) and relative to head perimeter (*EP*^*DV relative*^), respectively. **G)** Dorsal (blue) and ventral (red) eye perimeters relative to head perimeter (*EP*^*Dor relative*^, *EP*^*Ven relaive*^) across stages. **H-I)** Post-embryonic growth of the eye anteroposterior perimeter, expressed both in absolute terms (*EP*^*AP*^) and relative to head length (*EP*^*AP relative*^), respectively. **J)** Anterior (blue) and posterior (red) eye perimeters relative to head length (*EP* ^*Ant relative*^, *EP*^*Post relaive*^) across stages. **K)** Head perimeter (*HP*) and head length (*HL*) across stages. **L-M)** Number and facet diameter (*FD*) of ommatidia across stages, respectively. Sample sizes: *E-G, K*_*Head perimeter*_: *n*=3-5 per stage; *H-K*_*Head length*_: *n*=2-4 per stage; *L-M*: *n*=2-5 per stage. In *E-M*, solid lines represent the results of the regression analyses, performed either in logarithmic (*E, H, K, L*) or natural (*F, G, I, J, M*) scale. Regression slope estimates and significance levels are reported in Table S1. Insect stage was used as a numerical variable in the regression analyses shown, including the adult stage as the sixth stage. Filled circular markers with error bars represent the mean values estimated by the regression models, while semi-transparent markers show individual data points (raw data) for each insect. Individual identity is coded by the specific combination of shape and fill in the semi-transparent markers. Filled square markers with error bars represent observed mean values for the adult stage, while crosses represent the estimated mean values for the adult stage based on an extrapolation of the growth trend of the nymphal instars. In all cases, error bars denote standard errors. Asterisks represent significant differences (p-value <0.05) between estimated and observed values of the adult stage. When two eye perimeters were plotted in the same figure, a paired t-test was also performed to assess differences between the observed values of the adult for both perimeters (i.e. dorsal vs. ventral, anterior vs. posterior). In this cases, if differences were significant, a black asterisk was also added. Test statistics (t) and degrees of freedom (df) for the t-tests are reported in Table S2.

We performed a quantitative analysis of the eye size across stages for its anteroposterior and dorsoventral extents (Fig. 2 A-D). This analysis reveals that the eye dorsoventral perimeter (*EP*^*DV*^) in the adult stage is significantly larger than expected if we assume an exponential growth of this trait throughout stages (Fig. 2E; 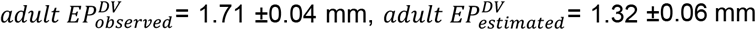, t-test: p< 0.001, see Tables S1 and S2 for more statistical details). Notably, a deviation from the growth trend in the adult stage also occurs for head perimeter (*HP*), but with opposite sign: while eye perimeter accelerates its growth rate in the last stage, head perimeter decelerates its own, resulting in a smaller head perimeter than expected for the adult (Fig. 2K; 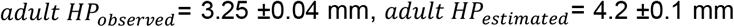, t-test: p< 0.001). For this reason, eye perimeter relative to head perimeter has a more dramatic change between expected and observed values in this last stage (Fig. 2F; 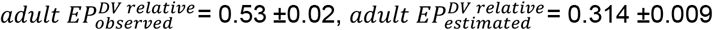, t-test: p< 0.001).

In addition, we separately analyzed the development of the eye dorsal and ventral perimeters relative to the head perimeter (Fig. 2G). This analysis shows that the deviation from the growth trend in the adult stage is significant for both eye perimeters (Fig. 2G; 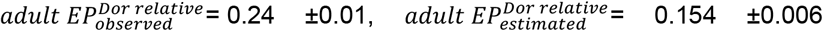 t-test: p^Dor^< 0.001, blue asterisk; 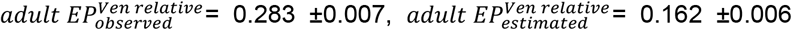 t-test: p^Ven^< 0.001, red asterisk). Finally, we observed that, across nymphal instars, there are no differences between the ventral and dorsal perimeters; whereas in the adult, the ventral perimeter is larger than the dorsal one resulting in an asymmetrical eye (t-test: p^DorVen^= 0.004, black asterisk).

With respect to the anteroposterior perimeter of the eye (*EP*^*AP*^), a shift in the development trend of the eye is also evinced for the adult stage. If we assume an exponential growth throughout stages, the eye perimeter observed for the adult is significantly larger than expected (Fig. 2H; 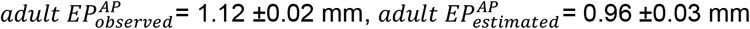, t-test: p= 0.005). The growth trend of head length (*HL*) also shows a deviation for the adult stage, but in the opposite direction: the observed value for the adult is significantly smaller than expected (Fig. 2K; 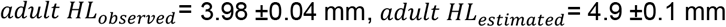, t-test: p< 0.001). Because of this, the deviation in the growth trend is much greater between expected and observed values for eye perimeter relative to head length (Fig 2I; 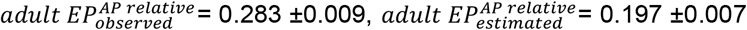, t-test: p= 0.002).

Additionally, we separately analyzed the development of the eye anterior and posterior perimeters relative to the head perimeter. This analysis shows that the deviation from the growth trend in the adult stage is significant for both perimeters of the eye (Fig. 2J; 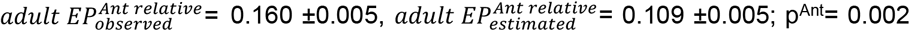, blue asterisk; 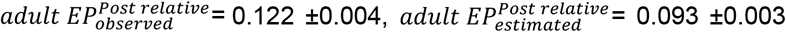 t-test: p^Post^= 0.004, red asterisk). Finally, we observed that the anterior region of the eye is larger than the posterior one in both nymphal and adult stages (t-test: p^AntPost^ < 0.001, black asterisk).

The analysis of the number of ommatidia (NO) along ontogeny revealed an exponential growth for this trait throughout all the developmental stages, *i*.*e*., no deviation was observed for the adult in this case (Fig. 2L). However, a trend deviation in the adult stage was observed for the growth of facet diameter (FD) across stages (Fig 2M; *adult FD*_*observed*_ = 57.7 ±1.2 µm, *adult FD*_*estimated*_ = 47.9 ±0.9 µm, t-test: p= 0.001).

Given that the main differences in eye growth were found between the last (fifth) nymphal instar and the adult stage for *R. prolixus*, the following analyses will focus exclusively on these two stages.

### 3.2 Fluorescent pseudopupil

The most traditional method for describing the functional morphology of compound eyes involves studying the principal pseudopupil to determine the optical axes of the ommatidia (42). In bright-pigmented eyes, the pseudopupil appears as a dark spot, revealing the ommatidia pointing towards the viewer. However, as wild-type *R. prolixus* has dark-pigmented eyes, a dark pseudopupil cannot be visualized. Notably, under an epifluorescence microscope, a bright pseudopupil can be revealed in dark-pigmented eyes due to photoreceptors' fluorescence (Fig. 3 A-B_i_; (43–45)). However, this phenomenon is visible only for a very brief period of time due to a light-adaptation process that involves the displacement of screening pigments (Fig. 3B_i-v_; (31, 33)). Because of the same adaptation process, antidromic illumination was neither appropriate for revealing a sustained pseudopupil. For this reason, in order to bypass the adaptation process, here we used red-eyed mutants, which do not possess the screening pigments involved in this mechanism (Fig. 3C; (34)). Thus, we could illuminate the eye for prolonged periods of time without any significant decrease in fluorescence making them suitable for imaging the pseudopupil (Fig. 3D). Additionally, in some preparations we intensified this fluorescence by injecting a fluorescent marker in the thorax of the animal to increase contrast (46).

**Figure 3.**
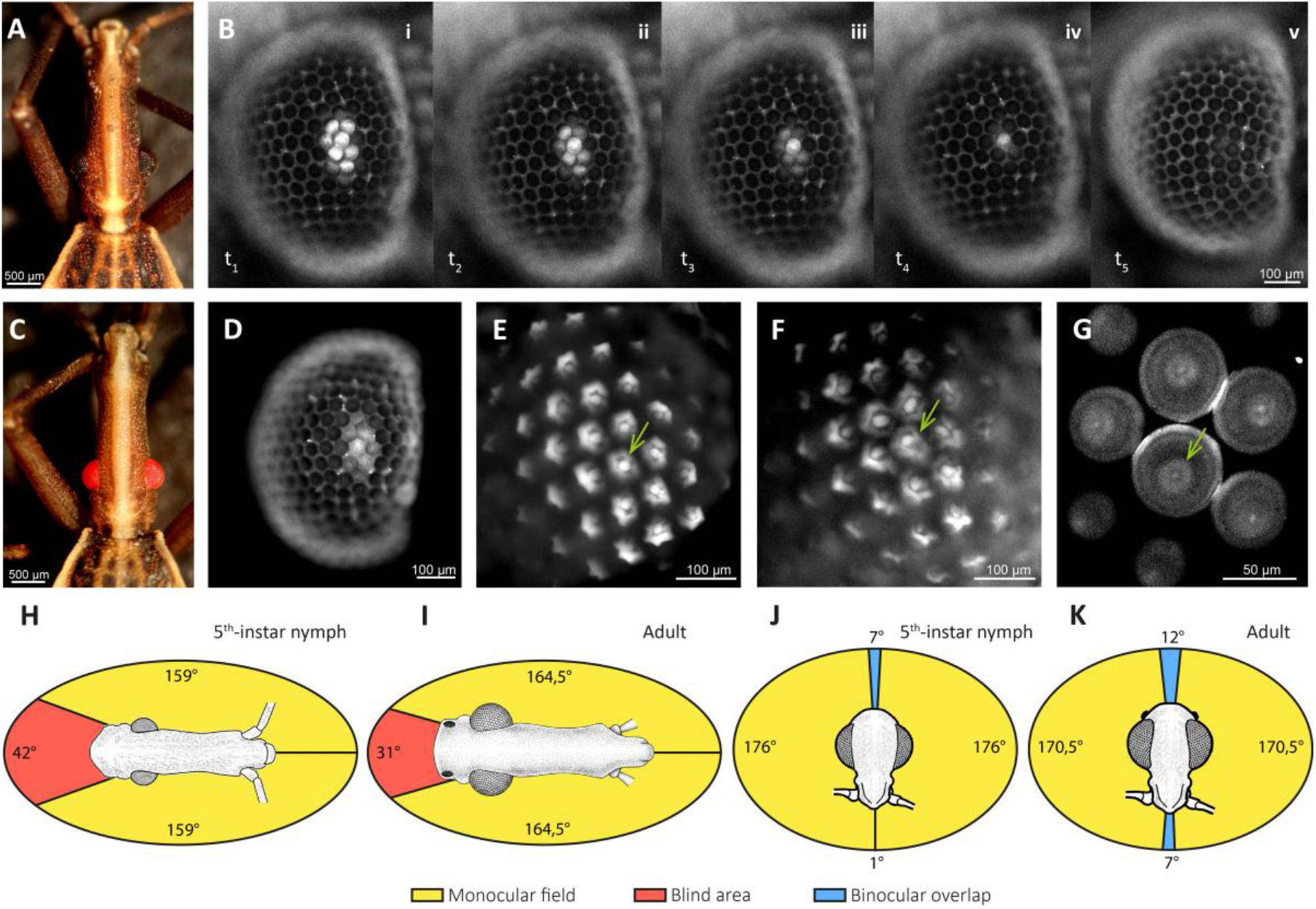
Autofluorescent pseudopupil and visual field of *R. prolixus*. **A, C)** Dorsal view of a black-eyed (wild-type) and a red-eyed (mutant) fifth-instar nymph, respectively. **B)** Time-lapse sequence (t1-t5) of a dark-pigmented eye showing the gradual fading of the pseudopupil in response to continuous focal illumination under an epifluorescence microscope. **D)** Lateral view of a red-pigmented compound eye under an epifluorescence microscope, showing a constantly bright pseudopupil (no fading). **E-F)** Open rhabdoms of photorreceptors of *R. prolixus* dark-pigmented and red-pigmented compound eyes, respectively, revealed via epifluorescence with water immersion. **G)** Open rhabdoms revealed by confocal microscopy. **H-I)** Anteroposterior visual field of a fifth-instar nymph and an adult, respectively (*n*=2 per instar). **J-K)** Dorsoventral visual field of a fifth-instar nymph and an adult, respectively (*n*=2 per instar). Green arrows in *E-G* indicate the position of an open-rhabdom, with its outer ring composed of photoreceptors 1-6 and central pair of photorreceptors 7-8. Scale bar in *B* applies to all images (*i-v*) in the row.

We have confirmed that in both eye phenotypes, the structures responsible for the autofluorescence phenomenon are the photoreceptors. In Figure 3 E-G (green arrows), the open rhabdoms typical of the heteropteran compound eye can be visualized (36). The open rhabdom consists of an outer ring composed of six rhabdomeres (R1-6) surrounding a central pair of rhabdomeres R7-8 (31, 33). The structure and shape of the open rhabdom seem to be well preserved in both the dark-eyed (Fig. 3E) and red-eyed insects (Fig. 3F). Additionally, we studied the visual defensive responses evoked in animals of both phenotypes when presented with a moving visual danger stimulus (Fig. S1). We found no differences between the responses of the two groups, suggesting that visual processing and sensitivity are not affected in red-eyed mutants. All these observations support the use of red-eyed mutants to study the functional morphology of the *R. prolixus* eye, as they indicate both structural and functional integrity.

### 3.3 Visual field

We determined the visual field of *R. prolixus* eyes by mapping the pseudopupil in the margins of the left and right eye in both the anteroposterior (horizontal) and dorsoventral (vertical) directions. Both fifth-instar nymphs and adults exhibit no binocular overlap in the horizontal plane, either in the frontal or rear visual fields (Fig. 3 H-I). While the frontal visual field is fully sampled in both stages, the posterior field is partially captured. The blind area corresponds approximately to the area subtended by the animal’s own body. The overall visual field in the horizontal plane is similar across both stages, with horizontal fields of view of 318° and 329° for the fifth-instar nymph and the adult, respectively. In contrast, the vertical visual field extends across a full 360° range and exhibits slight dorsal and ventral overlap in both stages (Fig. 3 J-K). These overlaps are slightly larger in the adult than in the nymph (dorsal overlap: nymph 7°, adult 12°; ventral overlap: nymph 1°; adult 7°).

### 3.4 Visual acuity and optical sensitivity

The interommatidial angles were determined by identifying the angular axes of the ommatidia through pseudopupil mapping across a horizontal and a vertical transect of the eye. Different shapes and sizes of the fluorescent pseudopupil in three angular positions along these two transects in an adult eye are shown in Figures 4A and D. A larger pseudopupil implies smaller interommatidial angles and, consequently, greater sampling resolution in that region of the eye. This is evident in the horizontal transect when comparing the pseudopupil horizontal dimension in the posterior (Fig. 4A_i_) and anterior regions of the eye (Fig. 4A_iii_): the latter exhibits a pseudopupil that recruits more ommatidia than the former (i.e. the latter possesses smaller interommatial angles and higher sampling resolution). A quantitative analysis along the horizontal transect shows that, for both stages, the interommatidial angle decreases towards the anterior region of the eye, resulting in approximately a 2.3-fold increase in sampling resolution (47, 48) in the frontal visual field (Fig. 4B; comparison between –32° and +82° in the adult, and between -21° and +64° in the nymph). In contrast, facet diameters shows a large increase in the adult with respect to the nymph (Fig. 4C). This difference in facet diameters is rather constant along the transect. The average facet size in the adult (65.3 ±0.5 µm) increases 62% relative to the nymph (40.4 ±1.6 µm), which implies a significant increase in the eye optical sensitivity.

**Figure 4.**
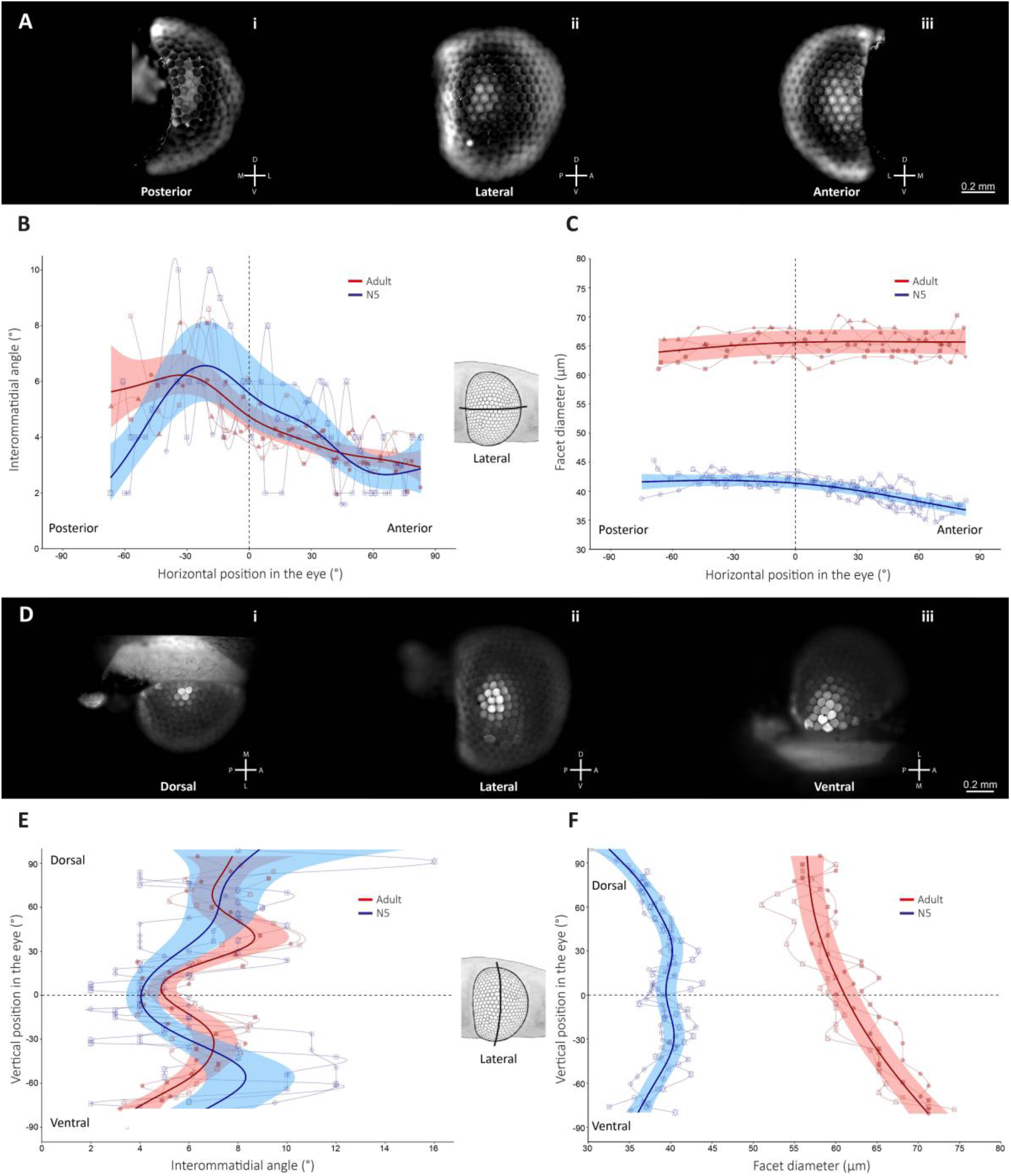
Sampling resolution and optical sensitivity of the *R. prolixus* compound eye. **A)** Views of the fluorescent pseudopupil at different regions along the horizontal (anteroposterior) transect of the eye: **i)** posterior; **ii)** equatorial; **iii)** anterior. **B-C)** Interommatidial angles (°) and facet diameters (µm) along the horizontal transect of the eye (°) for the fifth-instar nymph (blue) and the adult (red), respectively. **D)** Views of the fluorescent pseudopupil at different regions along the vertical (dorsoventral) transect of the eye: **i)** dorsal; **ii)** equatorial; **iii)** ventral. **E-F)** Interommatidial angles (°) and facet diameters (µm) along the vertical transect of the eye (°) for the fifth-instar nymph (blue) and the adult (red), respectively. Solid lines with shaded bands represent mean values with 95% confidence intervals. The curves result from fitting generalized additive mixed models (GAMMs) to the data from all eyes measured for each stage. Thin solid lines represent individual traces of each eye analyzed, and result from applying a monotone Hermite spline interpolation (following the method of Fritsch and Carlson) to data of each individual eye, for illustrative purposes only. Individual identity is coded by the specific combination of shape and fill in the semi-transparent markers. Anatomical axes are shown in *A* and *D* for head orientations (D: dorsal; V: ventral; M: Medial; L: Lateral; A: anterior; P: posterior). Illustrations of head sections between *B-*C and *E-F* indicate the locations of the transects sampled in the eye. Scale bars in *A* and *D* apply to all images (*i-iii*) within each respective row.

Along the vertical transect of the eye, variations in the interommatidial angles were less smooth than along the horizontal transect (Fig. 4E). For both stages, we found large interommatidial angles in the dorsal region of the eye, a local minimum at the eye equator and low angles towards the ventral region of the eye. Only in the ventral-most region of the eye did we observe a slight reduction in the interommatidial angles of the adult eye relative to the nymph eye. The facet diameters were consistently larger in the adult compared to the nymph along the entire transect (Fig. 4F). As observed in the horizontal transect, the average facet diameter is approximately 60% larger in the adult (61.6 ±4.5 µm) relative to the nymph (38.5 ±2.0 µm). However, whereas in the nymph the largest ommatidia are located at the eye equator and decrease in size in the dorsal and ventral directions, in the adult, ommatidial size increases from the equator towards the ventral region, reaching its maximum at the ventral-most region of the eye. These facet diameter profiles indicate that optical sensitivity peaks at the eye equator in the nymph, whereas in the adult, peak sensitivity shifts to the ventral-most region of the eye.

## Discussion

Here we have characterized the functional morphology of the compound eye of the kissing bug *Rhodnius prolixus* and the changes it experiences along ontogeny. These eyes are approximately hemispherical-shaped eyes which allow a whole visual sampling of the surrounding space with the exception of the space region occupied by its own body (Fig. 1). We have observed that in the adult stage the eye experiences a growth that exceeds the increment expected by extrapolating the growth rate across nymphal instars (Fig. 2). The size increment occurs in both the anteroposterior and dorsoventral perimeters of the eye (Fig. 2E and H, respectively), as well as in these perimeters normalized to head size (Fig. 2F and I). These increases in size are more pronounced in the anterior and ventral regions of the eye (Fig. 2G and J). Regarding eye sampling resolution, we observed no substantial differences between the last nymphal instar and the adult stage, either along the eye equator or along the vertical transect in the lateral region of the eye (Fig. 3B and E). The only exception was a slight increase in resolution in the ventral region of the adult eye. The ommatidial diameter, instead, was found to be considerably larger than expected in the adult stage. Along the eye equator, this parameter was rather constant in both the last nymphal instar and the adult stage (Fig. 4C). In contrast, along the vertical transect, ommatidial diameter reaches its maximum at the eye equator for the fifth-instar nymph, while in the adult stage the largest ommatidia are found in the ventral region (Fig. 4F). Thus, the area of highest sensitivity along the vertical transect shifts from the equatorial region in the nymphal instar to the ventral region in the adult.

Based on our observations in *R. prolixus*, we analyzed how the compound eyes of other triatomine species are modified throughout ontogeny. We found that *R. prolixus* is not the only triatomine species that shows a considerable enlargement of the eyes in the adult stage. *Triatoma infestans*, for example, also exhibits a significant enlargement of the eye, which is also more pronounced in the anterior and ventral regions (Figs. 5A and S2). In addition, although we were unable to quantify the eye size of the nymphal and adult stages in insects of the genus *Panstrongylus*, bibliographic and online photographic records show that the nymphal stages of this genus possess small compound eyes, whereas adults possess large eyes (49, 50). Figure 5B presents photographs from different perspectives of the *P. megistus* eye, illustrating the eye growth between the last nymphal instar and the adult stage. A particular condition regarding adult eye size is observed in animals from the genus *Mepraia*. In the last nymphal instar, animals possess relatively small eyes (49); however, in the adult stage the presence of a large eye is contingent upon the presence of wings. Typically, adult females of this genus are apterous (wingless) and possess relatively small compound eyes, whereas males develop wings and exhibit significantly larger eyes (Fig. 5C; (51)). Furthermore, in *M. spinolai*, there are also apterous males. Interestingly, the eye size in these males is relatively small, as observed in females (Fig. 5C). In summary, animals of the genus *Mepraia* also exhibit eye enlargement in the adult stage, which is closely linked to wing development. Thus, in all specimens from the genera *Rhodnius, Triatoma, Panstrongylus* and *Mepraia* that we could analyze, the development of large compound eyes in the adult is tightly associated with the presence of wings.

**Figure 5.**
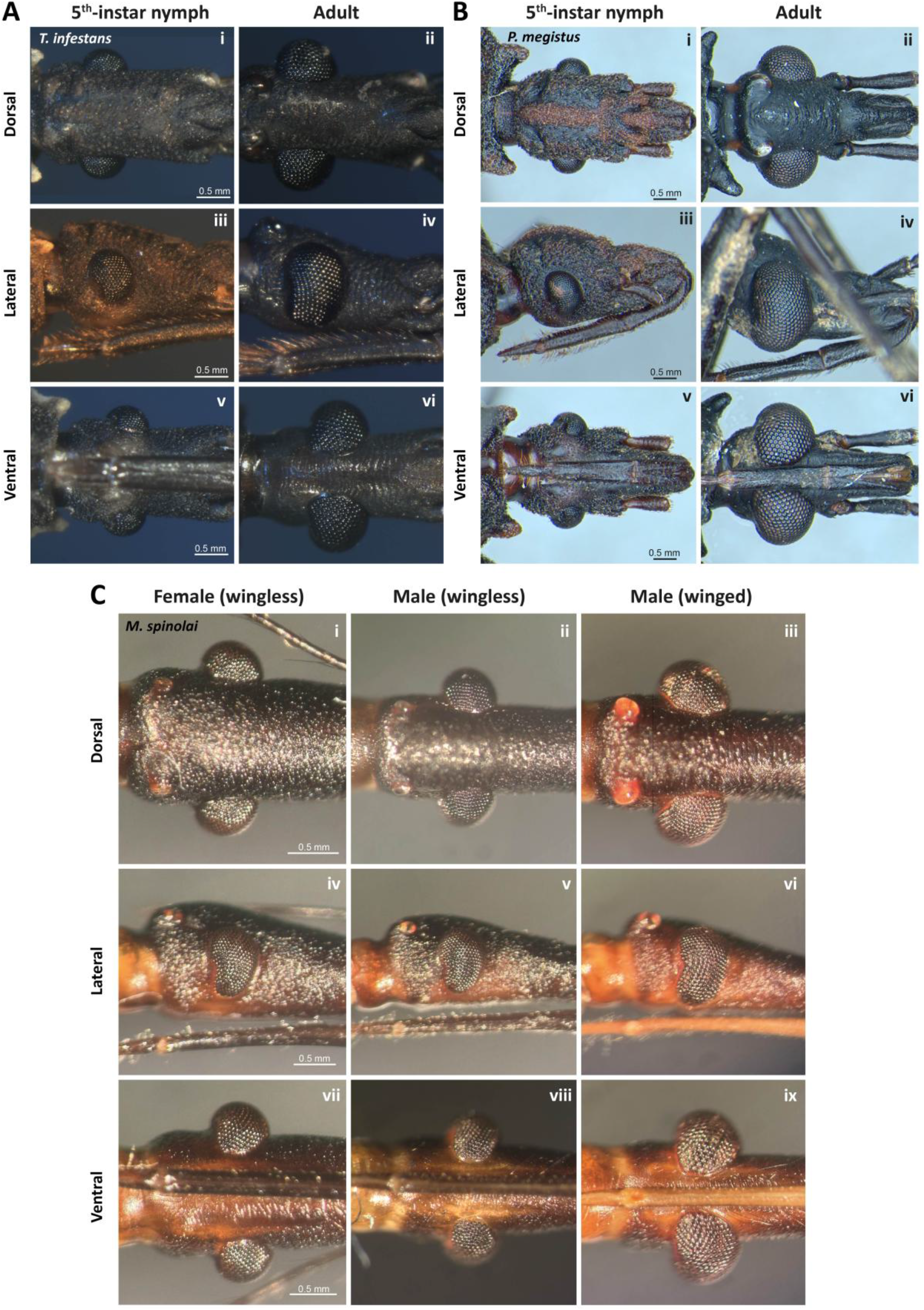
External morphology of the compound eyes of different triatomine species. **A-B)** Different views of the heads of a fifth-instar nymph and an adult of *Triatoma infestans* and *Panstrongylus megistus*, respectively: **i-ii)** dorsal; **iii-iv)** lateral; **v-vi)** ventral. **C)** Different views of the heads of a wingless adult female, a wingless adult male and a winged adult male of *Mepraia spinolai*: **i-iii)** dorsal; **iv-vi)** lateral; **vii-ix)** ventral. Scale bars in *A-C* apply to all images (*i-ii*) within each respective row.

The analyses of the eye sampling resolution performed in *R. prolixus* so far do not reveal a particular area of the eye in which resolution changes significantly between the last nymphal instar and the adult stage. As a distinctive characteristic of adult eyes, we observed a considerable ventral enlargement with respect to the eye of the last nymphal instar. Although in the adult stage the eye grows in both dorsal and ventral directions, it extends ventrally until nearly contacting the contralateral eye at the head midline. Additionally, whereas in the last nymphal instar the ommatidial size is dorsoventrally symmetrical (light blue curve in Fig. 4F), in the adult, ommatidial size increases monotonically from the dorsal to the ventral region of the eye, where it reaches its maximum (red curve in Fig. 4F). Thus, in *R. prolixus*, the adult eye becomes asymmetrical in both shape and ommatidial size. We could directly confirm that the eye transitions from being symmetrical in the last nymphal instar to asymmetrical in the adults of *T. infestans, P. megistus*, and winged adult males of *M. spinolai* (Fig. 5). In addition, we analyzed the evidence available in the bibliography and in online photographic records to evaluate adult eye symmetry (Table S3; 49, 50). After visually inspecting 39 species from 10 genera, we observed that in 36 species from 9 genera the adult eye possesses a greater ventral than dorsal extension and larger ommatidia in the ventral than in the dorsal region (Table S3). Thus, an asymmetrical adult eye appears to be the most likely eye pattern among triatomines.

As mentioned before, triatomines are obligate hematophagus insects that feed on warm-blooded hosts almost exclusively during the night. Throughout their lives, kissing bugs naturally seek to hide in folds or dark crevices that serve as shelters from which they perform foraging excursions. Olfactory and thermal cues have been considered the main sensory drivers of feeding behavior (52). A role of vision in this behavior has been classically neglected (discussed in (27)). We have recently shown that vision in triatomines plays a clear role in supporting defensive behaviors. Directional escaping, freezing or combination of these responses can be visually evoked in a fast and flexible manner according to the evaluation of the impending risk that the visual stimuli entail. Laterally placed eyes possessing a panoramic view of the world with a minimal posterior blind spot are characteristic of prey animals (53). Such orientation of the eyes allows scanning almost the 360° horizon around them, constantly on the lookout for eventual threats, for example, on foraging excursions. Thus, the visual field outlined here for the *R. prolixus* eye are consistent with previous finding indicating a role of the visual system in supporting defensive behaviors (27).

Adult triatomines become exposed to predators not only in foraging excursions but also in dispersal flights. Therefore, the great enlargement of the eye in the adults could serve to better cope with the additional risk that dispersal flights entail. This hypothesis is supported by the fact that large eyes are found only in triatomines that possess wings. Furthermore, the ventral expansion of the eye in winged triatomines could be associated with the novel features of the aerial niche. In contrast to terrestrial locomotion, flight exposes the insect to threats approaching from below. A more developed ventral region of the eye could address the need for monitoring the space beneath the animal. Additionally, stimuli approaching from ventral directions may exhibit lower contrast against the ground than those approaching from above, which are typically silhouetted against the sky. This could explain why the area of ventral ommatidia is approximately twice the area of the dorsal ones, providing greater light sensitivity.

In addition to supporting defensive behaviors, vision has been shown to be involved in dispersal flights. Dispersal flights generally occur in starving adults that do not have access to blood sources (11). Under controlled laboratory settings, it was observed that take-off activity is oriented towards focal sources of light (19, 22). In the field, the light of houses, the light of streets as well as traplights in different natural environments have been shown to attract dispersing triatomines (e.g. (20, 23, 24), respectively). These directional light-sensitive behaviors may benefit significantly from a sensitive compound eye. Similarly, flight control could also benefit from it. Regarding the ventral development of the eye, it has been shown that visual motion flow in the ventral visual field of different insects can assist in controlling flight speed, altitude, and landing maneuvers (e.g. (54–56), respectively). In addition, information on light polarization in this region of the visual field helps various arthropods orient their flight direction (57). However, despite the epidemiological importance of triatomine dispersal flights, studies characterizing these flights or their control have not yet been conducted. Therefore, it is currently difficult to discuss the feasibility of these hypothetical visual functions.

Kissing bugs have been intensively studied, partly due to their particular feeding habit, but mainly because these insects are the vectors of Chagas disease, the most important endemic disease from South to North America expanding now to other continents in the world (58). Our current findings reveal that eye size in four triatomine genera - including the genera of the primary vectors of Chagas disease in South and Central America (*T. infestans, R. prolixus, Triatoma dimidiata, P. megistus*; (59)) - is larger than predicted by the nymphal eye growth rate. This enlargement of the eye is significantly more pronounced in the ventral than in the dorsal region, a pattern we also identified in several other species of these and other genera. The ubiquity of this dorsoventral asymmetry suggests it is a basal trait within the group. While the role of vision in triatomine behavior has been historically overlooked in relation to other sensory systems, the characterization of the compound eye outlined here provides a solid framework for testing specific hypotheses regarding the visual ecology of adult triatomines.

## Materials and Methods

### 2.1 Animals

*R. prolixus* (Stäl 1859) was reared at the *Bioterio de Animales No Tradicionales* of the *Facultad de Ciencias Exactas y Naturales* (*Universidad de Buenos Aires)*. All stages (egg, nymph with its five different instars and adult) were kept in a rearing chamber at 28 ± 1ºC, 30-60% relative humidity with an inverted 12L:12D photoperiod (scotophase starting at 8:00 h and photophase at 20:00 h). Once a week, all individuals were fed *ad libitum* on live hens (handled according to the biosafety rules from the *Servicio de Higiene y Seguridad* of the university) and grouped according to their developmental stage. All individuals used in this work were aged at least 7 days post-ecdysis. Both females and males were included in this work, but, as no differences were found between them, we have grouped them with no gender specification along the article.

Wild-type (black-eyed) individuals were used to study eye growth (studies presented in Figs. 1-2). Instead, red-eyed mutant animals were used for the studies of visual fields, sampling resolution and optical sensitivity (studies presented in Figs. 3-4). These mutants were chosen because of the advantages they possess for pseudopupil imaging (see Section 2.3).

To evaluate the visual performance of red-eyed mutants, we compared the responses to a moving visual threat between red-eyed and wild-type fifth-instar nymphs (Fig. S1). This instar was chosen because the insects are large enough to be handled easily and have no ocelli (only adults have), enabling us to limit the visual input for the studied evoked behaviors to the compound eyes (27). The behavioral experiments were conducted during the scotophase, the time period in which these insects exhibit their maximal locomotor activity and display most of their behaviors in nature (13, 14). Starved nymphs were transported to the experimental room 1-2 hours before the experiments began.

### 2.2 Morphometrics of compound eyes

To characterize the development of the compound eyes in *R. prolixus*, we acquired images of the head and eyes across all developmental stages using different imaging techniques, and then quantified various morphological variables: head length, head perimeter, dorsoventral and anteroposterior perimeters of the compound eyes, number of ommatidia and facet diameters (see below).

#### 2.2.1 Image acquisition

Fluorescence microscopy: This optical method was used to visualize eye growth because the autofluorescence of the cuticle of these insects yields excellent contrast of the compound eyes. Thus, to illustrate eye growth across stages (Fig. 1 G-I), images of *R. prolixus* heads in dorsal, ventral and lateral views were captured using an epifluorescence microscope (BX51WI, Olympus Corporation, Tokyo, Japan). The system was equipped with a 10X objective (NA 0.20) and a filter set consisting of a 525 nm dichroic filter, a 470-490 nm band-pass excitation filter and a long-pass 530 nm emission filter. The LED light source used (peak wavelength: 470 nm; Tolket, Buenos Aires, Argentina) was coaxially aligned with the imaging pathway of a CMOS camera (C11440, Hamamatsu Photonics, Hamamatsu, Japan).

Stereomicroscopy: We used stereomicroscope images of the heads in dorsal view and head cross-sections to quantify several head and eye morphological parameters (see Section 2.2.2). The images were acquired with a color CMOS camera (BFS-U3-63S4C-C, FLIR Integrated Imaging Solutions, Canada) attached to the stereomicroscope (SZX10, Olympus Corporation, Tokyo, Japan).

Confocal microscopy: This method enabled us to perform three-dimensional (3D) reconstructions of the eye, allowing us to count the total number of ommatidia and measure facet diameters (see Section 2.2.2). Images of the compound eyes were taken in a confocal microscope (LSM 980 with Airyscan 2, ZEISS, Germany). Head and thorax of the five nymphal instars and the adult stage of *R. prolixus* were excised together from their bodies and mounted on a graphite stick in order to place them on the microscope and easily rotate them. Taking advantage of the autofluorescence of the cuticle, images were acquired at an emission wavelength of 509 nm after exciting the sample with a 488 nm laser (objective: 10x/NA 0.45 – pinhole size: 38 µm).

#### 2.2.2 Morphometric measurements

Quantitative analyses of the images were conducted using FIJI (60). To assess eye growth along the anteroposterior extent of the eye, images of the entire head in dorsal view were used to measure: head length (HL), total eye perimeter (EP^AP^), and eye anterior and posterior perimeters (EP^Ant^ and EP^Post^, respectively). To account for growth along the dorsoventral extent of the head, cross-section images were used to measure: head perimeter (HP), total eye perimeter (EP^DV^), and eye dorsal and ventral perimeters (EP^Dor^ and EP^Ven^, respectively). We have used head dimensions to normalize eye size parameters. Head dimensions were chosen to estimate body size throughout development, rather than variations in other body measurements that include the abdomen, because head measurements exhibit less variability than those including the extensible abdomen length (61). Our measure of head length spanned from the posterior margin of the cervix (at the junction with the prothorax) to the base of the rostrum (Fig 2A). As *R. prolixus* head has a cylindrical section, the head perimeter in cross-section was simply estimated by fitting a circle that included both the dorsal and ventral region of the head not occupied by the eyes (Fig. 2C).

To obtain the cross-section of the head, after washing the sample in 96% ethanol for several minutes to soften the cuticle (61), we performed a clean transversal cut through the middle of the eyes. Head cross-sections were only performed on the last three nymphal instars and the adult stage, because in smaller animals we found difficulties to cut the heads without distorting the eyes. The analysis of the last three nymphal instars proved sufficient to characterize the eye development process. This approach was validated with data from other eye variables, for which regression models including all five nymphal instars yielded results consistent with those obtained from models restricted to the last three instars (data not shown).

To count the total number of ommatidia in the compound eye for each of the five nymphal instars and the adult stage of *R. prolixus*, we generated 3D representations of the eyes based on confocal Z-stacks. Additionally, for each reconstructed eye, a set of 4 to 6 ommatidia from the lateral region at the eye’s equator was selected to quantify their diameters. To do so, a circle was fitted to the facet contour.

#### 2.2.3 Data analysis

Data analyses were conducted in the R environment (version 4.3.1 (62)). General linear models were fitted using the nymphal instar as a continuous explanatory variable. The individual identity was used as a random effect in the cases where multiple measurements were taken from the same insect (i.e. dorsoventral and anteroposterior eye perimeters, and facet diameters). For each response variable, linear regressions in both natural and logarithmic scale were performed (61, 63, 64). The better of these two fits was chosen based on the Akaike information criterion (AIC (65)). Regression slope estimates and significance levels are reported in Table S1. A one-sample t-test was performed to determine if the adult observed mean value was significantly different from the adult mean predicted by the nymphal regression model (61). When multiple measurements were obtained per adult insect, an individual mean was calculated for each insect to prevent pseudoreplication in the t-tests. Test statistics (t) and degrees of freedom (df) for the t-tests are reported in Table S2. In all cases, a level of significance of 0.05 was used. If no significant differences were found in the t-test, the observed values for the adult stage were then included in the regression analysis.

### 2.3 Functional morphology

To characterize the visual field, sampling resolution and optical sensitivity of the compound eye of fifth-instar nymphs and adults of *R. prolixus*, we imaged the eye pseudopupil, which allowed us to determine the optical axes of the ommatidia and measure their facet diameters (42). The compound eye of wild-type *R. prolixus* is typically dark-pigmented; therefore, a dark pseudopupil could not be visualized. However, we relied on two key factors to achieve a clear visualization of the pseudopupil: first, the photoreceptors of *R. prolixus* fluoresce under specific conditions, producing a bright pseudopupil; second, red-eyed mutants of this species lack pupillary screening pigments. Consequently, even under continuous illumination, their photorreceptor fluorescence remains clearly detectable. For this reason, we conducted the optical characterizations using red-eyed mutants. Sampling resolution and facet diameters were quantified along two eye transects (Fig. 3): a vertical (dorsoventral) transect traversing the lateral visual field, and a horizontal (anteroposterior) transect traversing the eye equator.

#### 2.3.1 Eye preparation

The head of either a fifth-instar nymph or an adult was severed and sealed with beeswax to prevent desiccation. The antennae and the proboscis were immobilized to ensure unobstructed optical access to the eye. The head was then mounted onto a graphite stick using beeswax: if the eye transect to study was vertical, the stick was attached to the posterior end of the head, parallel to the longitudinal head axis; otherwise, if it was the horizontal transect, the stick was fixed proximally to the antennae, perpendicular to the longitudinal head axis. Due to the plastic properties of the beeswax, the head alignment was then precisely adjusted under a stereomicroscope.

In order to increase the contrast of the bright pseudopupil in some individuals, a fluorescent dye was injected to the animal prior to head excision, following a modified version of the protocol designed by Gutierrez et al. (46). Briefly, the animal was immobilized on a slide with double-sided tape. A syringe with an angle of 30° with the long axis of the bug was inserted in the meso-thorax, and a constant pressure to the syringe plunge allowed a constant flux of a total 15-20 µl of a dye solution, which ended after a few minutes. The haemolymph flow through the dorsal vessel of the bug was sufficient to carry the dye to the cephalic capsule, thus bathing the optic lobes and the retina. After 30 minutes, the increase in fluorescence of the pseudopupil was notorious. The dye used was a purified TAT-GFP fusion protein dissolved in PBS buffer (0.80 mg ml^-1^), but the technique worked with Lucifer Yellow as well.

#### 2.3.2 Custom-built goniometer

Once the head (containing dye, when applicable) was mounted, the graphite stick was placed inside a 3D-printed holder fitted over the shaft of a wireless rotary encoder (PS-3220, PASCO Scientific, Roseville, USA). The encoder wheel was coupled to a stepper motor (28BYJ-48, China) via an elastic rubber band. This custom-built goniometer was controlled by an Arduino Uno board in a closed-loop manner. The position of the encoder (and thus, of the eye) could be easily set with a custom Python script by specifying either step-wise 2° rotations or specific absolute angular positions.

#### 2.3.3 Image acquisition

Images of the pseudopupil were captured using the epifluorescence microscope and the filter arrangement detailed in 2.2.1. The system is suited for imaging both autofluorescent and GFP-induced pseudopupils (Figs. 3 and 4).

The eye was rotated in discrete 2° steps, spanning the full length of the analyzed transects. At each step, an image of the pseudopupil was captured. At both extremes of the eye, the process ended when the pseudopupil was no longer visible.

#### 2.3.4 Pseudopupil mapping method

Once we have captured the images along an eye transect, we analyzed the distribution of the visual axes of the ommatidia using the traditional pseudopupil mapping method, similar to that used in previous studies ((46, 66–68), reviewed in (42)). This method allows for the estimation of the angle of separation between neighboring ommatidia (i.e. the interommatidial angle), which is a measurement of the optical spatial resolving power of a compound eye. The interommatidial angles were obtained by measuring the shift of the pseudopupil center (i.e. the number of ommatidia that it moves across) for a known rotation of the eye. Typically, the optical axis of an ommatidium within a transect was determined by identifying the angular position where its brightness was maximal. In subsequent images, we identified the next ommatidium in the transect with a clearly definable optical axis. The local interommatidial angle for that region was then calculated by dividing the rotation angle by the number of ommatidia spanned. Additionally, we measured the facet diameter of each of these ommatidia whose optical axes had been determined, as a proxy of their optical sensitivity (42). For the dorsoventral and anteroposterior transects, the angular reference (0°) was the angular position where the head was precisely aligned in a dorsal or anterior view, respectively. Finally, in an independent group of animals (two fifth-instar nymphs and two adults), we determined the visual fields of the right and left compound eyes and assessed the binocular overlap in the frontal, rear, dorsal and ventral regions. To this end, for each eye of an animal at each of these regions, we identified the maximum angular position where the pseudopupil was fully visible and distinguishable from the fluorescence of non-aligned ommatidia. The mean angular extent of each eye visual field was then calculated for each developmental stage. All image measurements were carried out with *FIJI*.

#### 2.3.5 Data analysis

Data analyses were also conducted in the *R* environment (version 4.3.1 (62)). Generalized additive mixed models (GAMM) were fitted to describe the variations of interommatidial angles and facet diameters as functions of the eye angular position along the two chosen transects. A Gamma distribution was assumed for the response variables. We performed four independent GAMMs, one for each combination of transect (vertical/horizontal) and stage (nymph/adult). Angular position was included as a fixed effect, while the identity of the compound eye was specified as a random effect. For adult-stage GAMMs, individual identity was also included as a random effect since both eyes were sampled in some individuals. GAMMs were fitted using the *mgcv* package (69) and model assumptions were evaluated using the *DHARMa* package (70).

## Acknowledgments

We deeply thank Raquel Aparecida Ferreira, Frederico Dutra Kirst and the Coleção de Vetores de Tripanosomatídeos (Instituto René Rachou, Fundação Oswaldo Cruz) for sharing with us photographs of several species of the genera *Panstrongylus, Triatoma* and *Rhodnius*. The photographs of *Panstrongylus megistus* were included in Figure 5.

