## Supplemental Figures and Tables for "Ontogenetic expansion and regionalization of the triatomine compound eye supports flight-related vision"

**This PDF file includes:**

Figures S1 to S2
Tables S1 to S3
SI References

**Figures**

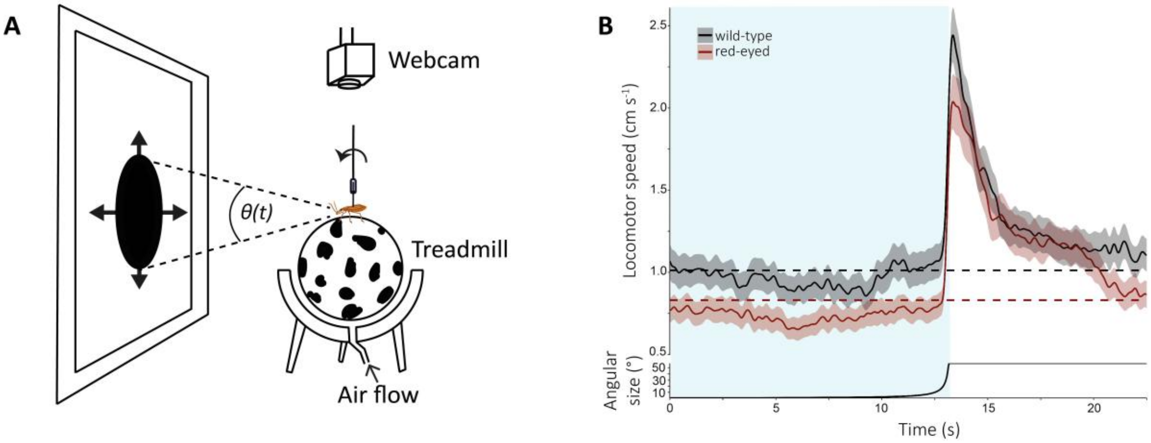


**Figure S1. Functional assessment of visual performance of red-eyed mutants using looming-evoked defensive behaviors.** The general experimental procedure followed the protocol described in Chialina[1] et al. 2025. **A)** Behavioral setup. A fifth-instar nymph was dorsally tethered to a freely rotating vertical rod and placed on an air-suspended black-spotted Styrofoam ball, where it could walk and rotate at will while keeping the same position in space. Videos were recorded with a webcam and then used to calculate the instantaneous speed of the animals at each frame with the software FicTrac[2]. The visual looming stimulus was programmed in Bonsai using the package Bonvision[3] and was presented on a screen located 20 cm away from the animal. It consisted of the simulation of a dark spherical object of 1 cm-diameter over a bright background, initially located at 100 cm from the animal, approaching at a constant velocity of 7.5 cm $s^{-1}$, reaching up to a distance of 1 cm from the animal. The initial diameter of the figure on the screen was 0.22 cm, subtending an angular size of 0.57° at the nymph's eyes. The final diameter of the figure was 23 cm, corresponding to an angular size of 57°. Every insect of each eye phenotype was presented with 5 presentations of the same looming stimulus. The intertrial interval was 30 s. Such intertrial interval yielded no habituation or sensitization of the defensive responses[1]. **B)** Average instantaneous speed (mean ± S.E.) of red-eyed mutants (red line, N=23) and wild-type insects (black line, N=22) when confronted with a visual looming stimulus. Light-blue shaded area highlights the time of stimulation. Dashed lines represent the mean spontaneous speed of wild-type (black line) and red-eyed (red line) insects during the 10 s prior to looming onset. Weber contrast of the stimulus was -0.99. Dynamics of the stimulus are shown at the bottom as the angular size of the simulated approaching object over time. To analyze statistical differences between the escape responses of the two phenotypes, we quantified the change in the distance traveled (DT) by the animals in response to the stimuli using smoothed data of the instantaneous speed. We calculated the area under the instantaneous speed trace from 1 s before to 3 s after the end of stimulus expansion. A general linear mixed model was fitted to analyze the distance traveled (DT) by the animals using the glmmTMB package[4]. The eye phenotype was used as a fixed effect. Model assumptions were evaluated using the DHARMa package[5]. We observed that the escape response evoked by a visual looming stimulus of red-eyed mutants were indistinguishable from that of dark-eyed insects: no statistical differences were found in the distance travelled (DT) between insects of the two eye phenotypes when faced with the same threatening stimulus (DTred-eyed= 5.240±0.494 cm, DTwild-type= 5.840±0.505 cm; $\chi$2= 0.716, df= 1, p= 0.397).

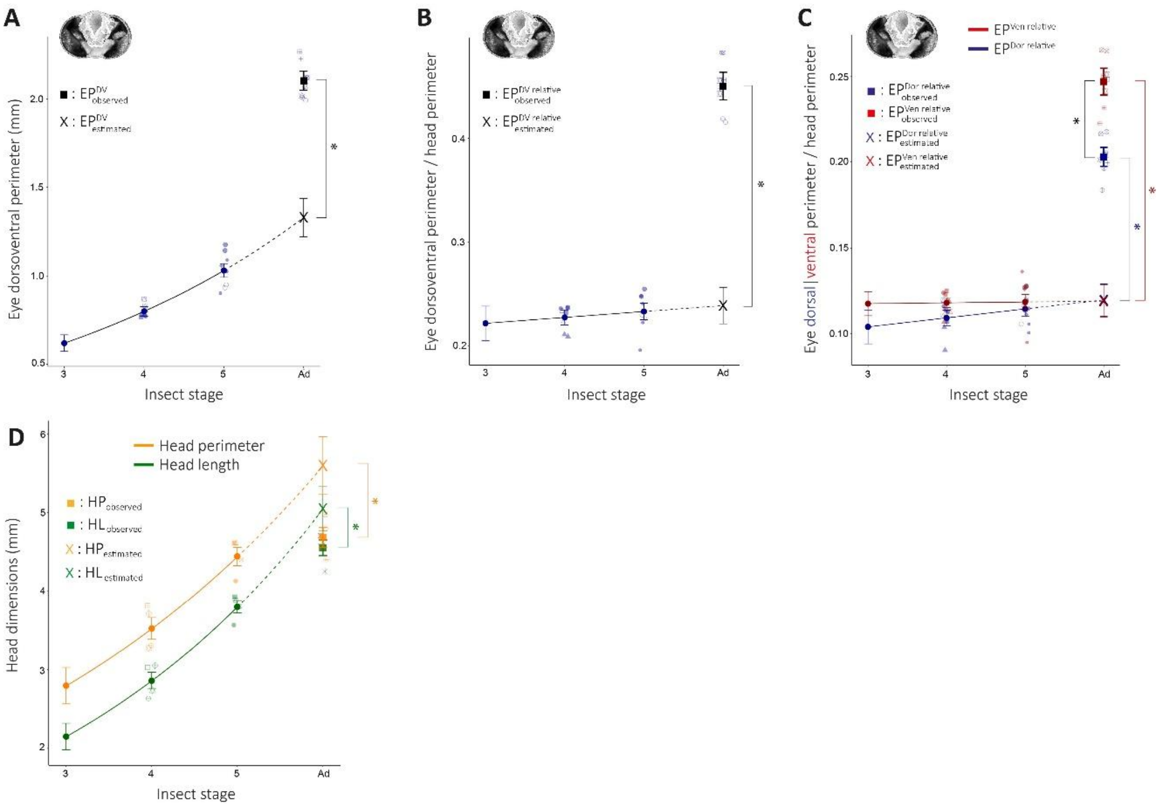


**Figure S2. Quantification of eye growth of *T. infestans*.** The protocol used to obtain these measurements and the regression analyses followed the same procedure as those performed for *R. prolixus* (see Materials and Methods section 2.2). **A-B)** Post-embryonic growth of the eye dorsoventral perimeter, expressed both in absolute terms ($EP^{DV}$) and relative to head perimeter ($EP^{DV\ relative}$), respectively. **C)** Dorsal (blue) and ventral (red) eye perimeters relative to head perimeter ($EP^{Dor\ relative}$, $EP^{Ven\ relaive}$) across stages. **D)** Head perimeter ($HP$) and head length ($HL$) across stages. Sample sizes: *A-D*: *n*=4-5 per stage. In *A-D*, solid lines represent the results of the regression analyses, performed either in logarithmic (*A, D*) or natural (*B, C*) scale. Insect stage was used as a numerical variable in the regression analyses shown, including the adult stage as the sixth stage. Filled circular markers with error bars represent the mean values estimated by the regression models, while semi-transparent markers represent observed individual data points (raw data) for each insect. Individual identity is coded by the specific combination of shape and fill in the semi-transparent markers. Filled square markers with error bars represent observed mean values for the adult stage, while crosses represent the estimated mean values for the adult stage based on the extrapolation of the growth trend of the nymphal instars. In all cases, error bars denote standard errors. Other conventions also as in Figure 2. T-tests:

$adult\ EP^{DV}_{observed}\ vs.\ adult\ EP^{DV}_{estimated}$ : t (df=3)= 18.2, p< 0.001;

$adult\ HP_{observed}\ vs.\ adult\ HP_{estimated}$: t (df=3)= -6.6, p= 0.007;

$adult\ EP^{DV\ relative}_{observed}\ vs.\ adult\ EP^{DV\ relative}_{estimated}$: t (df=3)= 15.9, p<0.001

$adult\ EP^{Dor\ relative}_{observed}\ vs.\ adult\ EP^{Dor\ relative}_{estimated}$: t (df=3)= 15.0, p< 0.001;

$adult\ EP^{Ven\ relative}_{observed}\ vs.\ adult\ EP^{Ven\ relative}_{estimated}$: t (df=3)= 16.4, p< 0.001 ;

$adult\ EP^{Dor\ relative}_{observed}\ vs\ adult\ EP^{Ven\ relative}_{observed}$: t(df=3)= -18.1, p< 0.001;

$adult\ HL_{observed}\ vs.\ adult\ HL_{estimated}$: t(df=3)= -4.6, p= 0.02.

### Tables

**Table S1.** Slopes and p-values for the regression models reported in Results 3.1. Unless otherwise specified, slopes are reported on a natural scale. $EP^{DV}$: Eye dorsoventral perimeter; $HP$: Head perimeter; $EP^{DV\ relative}$: Eye dorsoventral perimeter relative to head perimeter; $EP^{Dor\ relative}$: Eye dorsal perimeter relative to head perimeter; $EP^{Ven\ relative}$: Eye ventral perimeter relative to head perimeter; $EP^{AP}$: Eye anteroposterior perimeter; $HL$ : Head length; $EP^{AP\ relative}$: Eye anteroposterior perimeter relative to head length; $EP^{Ant\ relative}$: Eye anterior perimeter relative to head length; $EP^{Post\ relative}$: Eye posterior perimeter relative to head length; $NO$ : Number of ommatidia; $FD$: Facet diameter.

| Regression model | Slope | P-value |
|---|---|---|
| $EP^{DV}$ | Log-scale: 0.36 ±0.02 | $p< 0.001$ |
| $HP$ | Log-scale 0.30 ±0.02 | $p< 0.001$ |
| $EP^{DV\ relative}$ | 0.018 ±0.004 | $p< 0.001$ |
| $EP^{Dor\ relative}$ | 0.008 ±0.003 | $p= 0.007$ |
| $EP^{Ven\ relative}$ | 0.011 ±0.003 | $p< 0.001$ |
| $EP^{AP}$ | Log-scale: 0.31 ±0.01 | $p< 0.001$ |
| $HL$ | Log-scale: 0.319 ±0.007 | $p< 0.001$ |
| $EP^{AP\ relative}$ | -0.002 ±0.003 | $p= 0.6$ |
| $EP^{Ant\ relative}$ | 0.000 ±0.002 | $p= 0.9$ |
| $EP^{Post\ relative}$ | 0.000 ±0.001 | $p= 0.7$ |
| $Number\ of\ Ommatida\ (NO)$ | Log-scale: 0.426 ±0.004 | $p< 0.001$ |
| $Facet\ Diameter\ (FD)$ | 4.7 ±0.3 µm.instar$^{-1}$ | $p< 0.001$ |

**Table S2.** Test statistics (t) and degrees of freedom (df) for the t-tests reported in Results 3.1. *Observed*: observed mean value; *Estimated*: mean value estimated for the adult by the regression model.

| T-test | Statistic (t) | Degrees of freedom (df) |
|---|---|---|
| $adult\ EP_{observed}^{DV}\ vs.\ adult\ EP_{estimated}^{DV}$ | 11.5 | 4 |
| $adult\ HP_{observed}\ vs.\ adult\ HP_{estimated}$ | -19.3 | 4 |
| $adult\ EP_{observed}^{DV\ relative}\ vs\ adult\ EP_{estimated}^{DV\ relative}$ | 13.2 | 4 |
| $adult\ EP_{observed}^{Dor\ relative}\ vs\ adult\ EP_{estimated}^{Dor\ relative}$ | 9.1 | 4 |
| $adult\ EP_{observed}^{Ven\ relative}\ vs\ adult\ EP_{estimated}^{Ven\ relative}$ | 16.7 | 4 |
| $adult\ EP_{observed}^{Dor\ relative}\ vs\ adult\ EP_{observed}^{Ven\ relative}$ | -5.9 | 4 |
| $adult\ EP_{observed}^{AP}\ vs.\ adult\ EP_{estimated}^{AP}$ | 7.5 | 3 |
| $adult\ HL_{observed}\ vs.\ adult\ HL_{estimated}$ | -20.9 | 4 |
| $adult\ EP_{observed}^{AP\ relative}\ vs\ adult\ EP_{estimated}^{AP\ relative}$ | 9.8 | 3 |
| $adult\ EP_{observed}^{Ant\ relative}\ vs\ adult\ EP_{estimated}^{Ant\ relative}$ | 10.2 | 3 |
| $adult\ EP_{observed}^{Post\ relative}\ vs\ adult\ EP_{estimated}^{Post\ relative}$ | 7.8 | 3 |
| $adult\ EP_{observed}^{Ant\ relative}\ vs\ adult\ EP_{observed}^{Post\ relative}$ | 17.6 | 3 |
| $adult\ FD_{observed}\ vs.\ adult\ FD_{estimated}$ | 8.2 | 4 |

**Table S3.** Triatomine species assessed for the presence of dorsoventrally asymmetrical compound eyes in the adult stage. Sources: photographic records from the online collection of the Faculdade de Ciências Farmacêuticas de Araraquara[6] (A), and the following published material: Carcavallo et al. 1998[7] (B) and Paiva et al. 2021[8] (C).

| Species | Dorsoventral eye asymmetry | Source |
|---|---|---|
| ***Cavernicola*** *pilosa* | Yes | A |
| *C. lenti* | Yes | A |
| ***Dipetalogaster*** *maxima* | Moderate | A |
| ***Eratyrus*** *mucronatus* | Yes | A |
| ***Nesotriatoma*** *confusa* | Yes | A |
| ***Paratriatoma*** *leticularia* | Yes | A |
| *P. hirsuta* | Yes | B, C |
| ***Panstrongylus*** *lignarius* | Yes | A, B |
| *P. megistus* | Yes | A, B |
| *P. tibiamaculatus* | Yes | A |
| *P. rufotuberculatus* | Yes | B |
| *P. humeralis* | Yes | B |
| *P. herreri* | Yes | B |
| *P. chinai* | Yes | B |
| *P. geniculatus* | Yes | B |
| *P. lenti* | Yes | B |
| *P. tupynambai* | Yes | B |
| *P. lutzi* | Yes | B |
| *P. diasi* | Yes | B |
| *P. guentheri* | Yes | B |
| ***Psammolestes*** *arthuri* | Yes | B |
| *P. coreodes* | Yes | A |
| *P. tertius* | Yes | A |
| ***Triatoma*** *rubrovaria* | Yes | A |
| *T. sordida* | Yes | A |
| *T. platensis* | Yes | A |
| *T. brasiliensis* | Moderate | A |
| *T. pseudomaculata* | Yes | A |
| *T. vitticeps* | Yes | A |
| *T. nigromaculata* | Yes | A |
| ***Rhodnius*** *pictipes* | Yes | A |
| *R. pallescens* | Yes | A |

| | | |
|---|---|---|
| *R. neglectus* | Yes | A |
| *R. brethesi* | Yes | A |
| *R. neivai* | No | A |
| *R. ecuadoriensis* | Yes | A |
| *R. stali* | Yes | A |
| ***Mepraia** gajardoi* | Yes in males (brachypterous); No in females (apterous) | A |
| *M. parapatrica* | Yes in males (brachypterous or macropterous); No in females (apterous) | A |
